# Generation of a recombinant mimivirus by direct transfection of a PCR amplicon

**DOI:** 10.1101/2025.04.28.651011

**Authors:** Hiroyuki Hikida, Hiroyuki Ogata

## Abstract

Giant viruses are large double-stranded DNA viruses encoding hundreds of genes. These genes are considered to reprogram the host cell machinery, which, in turn, modulates the global nutrient cycles in the environment. However, the functions of giant-virus genes are largely unknown due to the lack of reverse-genetics systems. Recently, reverse-genetics systems have been developed in giant viruses infecting a free-living amoeba, *Acanthamoeba castellanii*. One of the methods relies on homologous recombination by plasmid transfection, but plasmid construction is generally laborious. In this study, we developed a method to generate a recombinant giant virus, mimivirus, with direct transfection of a PCR amplicon into host cells, a widely used approach in budding yeast and other organisms. Consequently, we successfully generated a recombinant mimivirus with a marker gene. Our results demonstrated that a bacterium-free method could be applied to giant viruses to generate recombinants, accelerating further experimental studies related to these viruses.

## Introduction

In general, viruses have small genomes, but some viruses, called giant viruses, have extremely large genomes, reaching 2.5 Mbp [1–3]. Their genomes encode various genes previously considered exclusive to cellular organisms, such as those related to translation, chromatin structure, and energy production [4–9]. These genes may modulate host metabolism to create favorable cellular environments for viral replication. In nature, the reprogramming of the host cellular machinery is considered to modulate global nutrient cycles [9,10]. Moreover, giant viruses encode various genes unique to viruses without homologous proteins in cellular organisms [11,12]. However, the functions of these giant-virus genes remain mostly elusive due to limitations in the reverse-genetics tools available to giant viruses and their host organisms.

*Acanthamoeba castellanii* is a free-living amoeba susceptible to diverse lineages of giant viruses. Since this amoeba has been studied for a long time, several molecular biology tools have been developed in this organism [13,14]. Therefore, the viral infection model using this amoeba is a favorable system for experimentally investigating giant viruses. Recent studies further established reverse-genetics systems in these amoeba-infecting giant viruses, paving the way to explore the gene functions of giant viruses [15–19]. One CRISPR-Cas9 system-based method allows for editing viral genome sequences, but this system can be applied only to those giant viruses that replicate inside the host nucleus [20]. Other giant viruses form their virus factory in the host cytoplasm, where the Cas9 protein fails to be recruited. For these giant viruses, homologous recombination using plasmid vectors is an alternative method to generate recombinant viruses [21,22]. However, plasmid construction is generally laborious.

Direct transformation of PCR amplicons is widely used to disrupt gene functions in the budding yeast *Saccharomyces cerevisiae* and is also used in other organisms [20–24]. Using this approach, we developed a method to generate recombinant giant viruses using PCR-based donor DNA. In this study, we focused on a mimivirus, an intensively studied giant virus. This virus harbors over one thousand genes, and homologous recombination is the only available tool for generating recombinants. We targeted the GMC-oxidoreductase 1 gene (previously known to be nonessential [21]) and successfully generated a deletion mutant. This result indicates that a PCR-based reverse-genetics system is also available in giant viruses, thereby potentially accelerating further genetic studies of giant viruses.

## Materials and Methods

### Cells and viruses

*Acanthamoeba castellanii* (Douglas) Page, strain Neff (ATCC 30010) was maintained with peptone-yeast extract-glucose (PYG) medium at 28°C. Acanthamoeba polyphaga mimivirus (APMV; NCBI Reference Sequence: NC_014649.1) was used as a mimivirus prototype [2,25,26].

### Plasmid construction

We constructed a plasmid that harbors the expression cassette of a marker gene with the APMV transcription machinery. A plasmid containing a fusion gene of a coding sequence of green fluorescent protein (GFP) and neomycin resistance (NEO) gene was purchased from VectorBuilder. APMV genomic DNA was extracted as described previously [27]. ORF of the GFP/NEO fusion protein was amplified from the plasmid using polymerase chain reaction (PCR) and the KOD One^®^ PCR Master Mix (Dye-free 2×PCR Master Mix) (KMM-101, TOYOBO, Japan). The upstream region of the APMV R659 gene and the downstream region of the APMV L274 gene were PCR-amplified using the APMV genomic DNA as a template. The amplified fragments were cloned into the backbone pBR322, which is linearized from the pUC19 vector using PCR, using In-Fusion^®^ Snap Assembly Master Mix (638948, TAKARA, Japan). The resultant plasmid was dubbed pMimV-KO-GFP/Neo. The primers used for PCR are listed in Table 1.

**Table 1.**
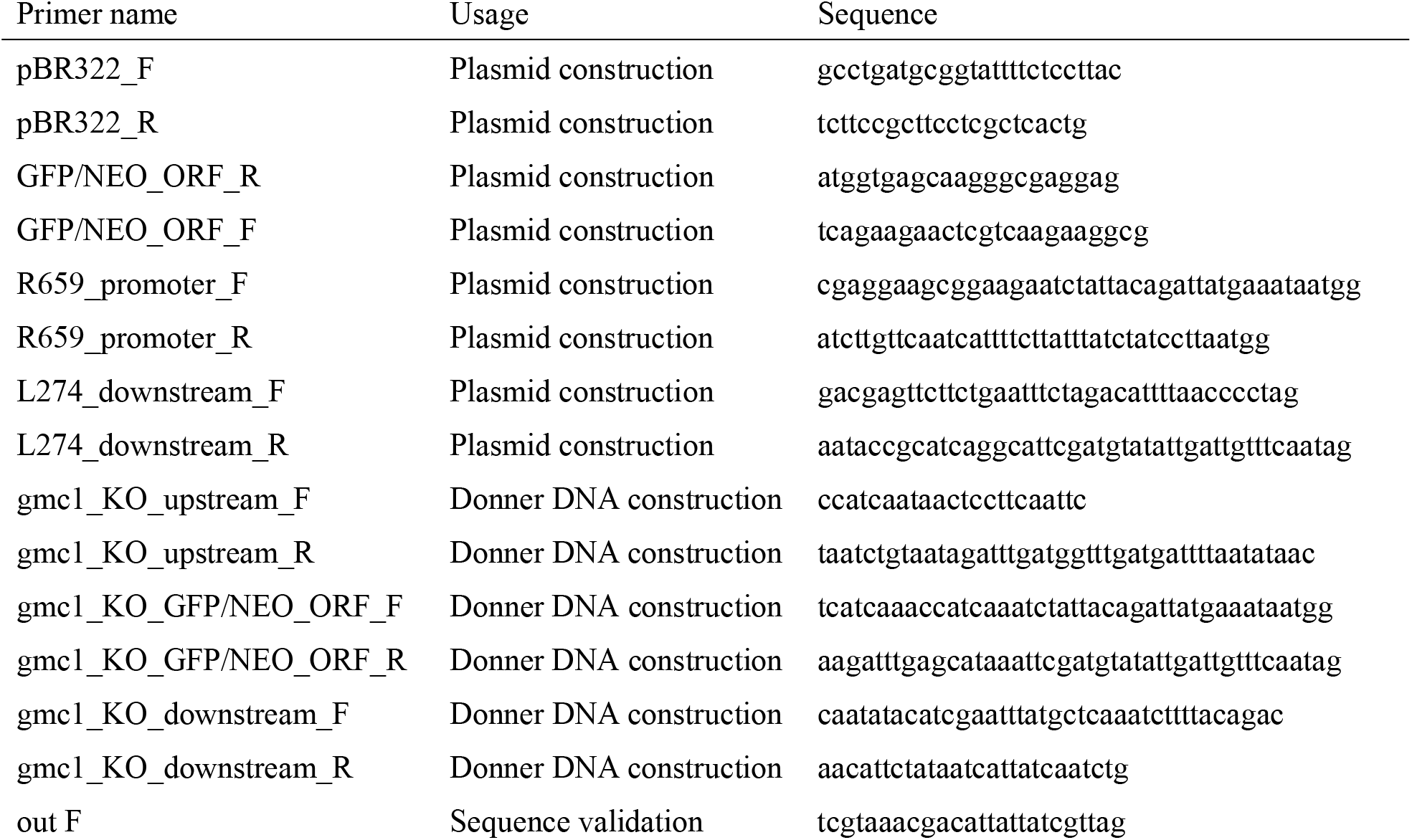
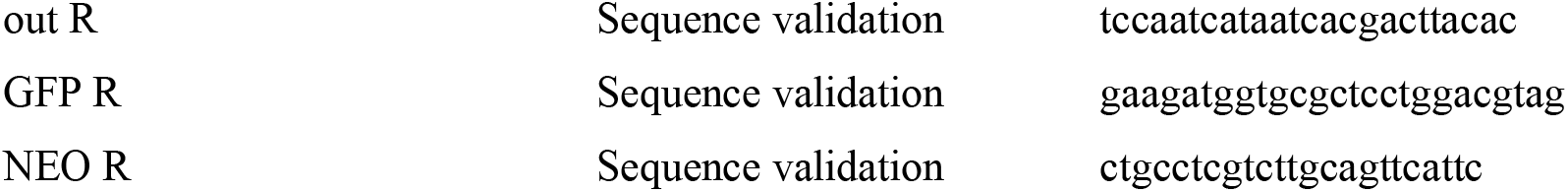
Primer List.

### Donor DNA construction

Donor DNA for homologous recombination was prepared as follows: the expression cassette for the GFP-NEO fusion gene and the upstream and downstream regions of the GMC-oxidoreductase 1 gene were PCR-amplified, followed by overlap extension using the KOD One^®^ PCR Master Mix in a single tube. The plasmid constructed above and the APMV genomic DNA were used as templates. Each primer concentration was 1.5 µM. The PCR condition was as follows: 25 cycles of 98°C for 10 s, 45°C for 5 s, and 68°C for 9 s, followed by 15 cycles of 98°C for 10 s, 55°C for 5 s, and 68°C for 5 s. The amplified DNA was separated using electrophoresis, and the area around the size of 3 kbp was cut, followed by gel extraction using the Monarch^®^ DNA Gel Extraction Kit (T1020, New England Biolabs). The extracted DNA was subjected to PCR using the KOD One^®^ PCR Master Mix and primers, gmc1_KO_upstream_F and gmc1_KO_upstream_R, and the amplicon size was validated by electrophoresis. The primers used for PCR are listed in Table 1. The PCR condition was as follows: 45 cycles of 98°C for 10 s, 45°C for 5 s, and 68°C for 15 s.

### Transfection

Amoeba cells at the growth phase were collected, and 5×10^5^ cells were seeded in a 35-cm Nunc EasYDish (150460, Thermo Scientific) with 2 mL of PYG medium. Twenty µL of the amplified PCR mix and 10 µL of PolyFect Transfection Reagent (301105, Qiagen) were added to 100 µL of sterilized phosphate-buffered saline. This transfection mix was incubated for 10 min at room temperature. Six hundred µL of PYG medium was added to this transfection mix, which was mixed gently by pipetting and supplemented into the dish. Immediately after the transfection, APMV was inoculated at a multiplicity of infection of 10, and the dish was incubated at 30°C. At 1-day post-infection, the supernatant was collected from the dish, and the remaining cells were removed by centrifugation at 2,000 rpm for 5 min at 4°C (Sorvall ST8FR, Thermo Scientific) and stored at 4°C as a viral suspension.

### Selection of recombinant viruses

In a 35-cm dish, 5×10^5^ cells were seeded with 2 mL PYG. The collected viral suspension was diluted ten times in PYG and inoculated into the amoeba. Subsequently, G-418 (16512-36, Nacalai Tesque, Japan) was added at a final concentration of 50 µg/mL and incubated at 30°C for 1 day. The supernatant was collected from the incubated dish, and the remaining cells were removed as described above. One hundred µL of the supernatant was mixed with 1×10^5^ amoeba cells in 10 mL of PYG medium and was dispensed to each well of a Nunc™ MicroWell™ 96-well plate (167008, Thermo Fisher). The plate was observed under a fluorescent microscope (CKX53 and U-LGPS, Olympus), and GFP-positive wells were collected. The collected viral suspension was further purified three times by end-point dilution, selecting GFP-positive wells.

### Confirmation of the recombination

The recombination was confirmed by PCR as follows. First, the purified recombinant virus was cultured in a 35-mm dish for 1 day. The supernatant was collected, and the remaining cells were removed, as described above. Next, the resultant supernatant was centrifuged at 15,000 rpm for 15 min at 4°C. The pellet of the viral particles was resuspended with 45 µL of 50 mM sodium hydroxide and incubated for 10 min at 95°C. Five µL of 1 M Tris-HCl at pH 8.0 was added to this reaction, followed by a dilution in 450 µL of Tris-EDTA buffer at pH 8.0. A part of the genomic sequence was PCR-amplified from the extracted DNA as described above and electroporated on a 1% agarose gel. The resulting gel was stained with Atlas ClearSight Gold DNA Stain (BioAtlas) for 10 min and imaged using FAS-BG LED BOX (Nippon genetics) and a 1 kb DNA Ladder PLUS (Nippon genetics) as the standard. The primers used for PCR are listed in Table 1. The PCR condition was as follows: 35 cycles of 98°C for 10 s, 45°C for 5 s, and 68°C for 20 s.

## Results and Discussion

We prepared the donor DNA for homologous recombination by overlap-extension PCR to concatenate the 500-bp flanking regions of the target gene (i.e., GMC-oxidoreductase 1) and an expression cassette of the marker gene (Fig. 1a). The expression cassette included the upstream of the APMV R659 gene and the downstream of the APMV L274 gene as a promoter and a 3’ untranslated region, respectively, because these two genes showed high expression from an early stage of infection [28]. To simplify the protocol, we amplified and concatenated the fragments in a single tube as a consecutive reaction (Fig. 1b). The resultant fragment was invisible in agarose-gel electrophoresis (Fig. 1c). However, after the gel extraction from the area around 3 kbp, we successfully PCR-amplified the resultant fragment (Fig. 1c).

**Figure 1.**
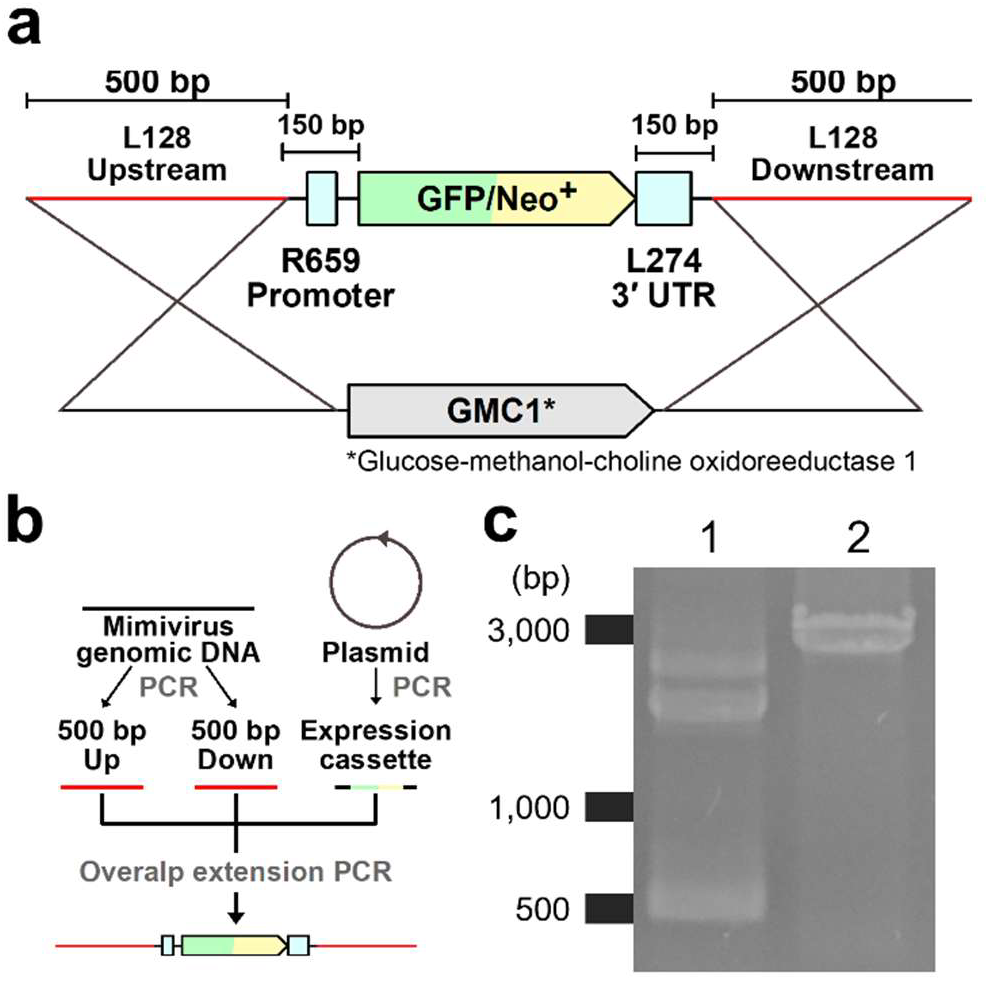
Donor DNA construction. (a) Schematic image of the donor DNA. (b) Schematic image of the overlap-extension PCR. The reaction was performed in a single tube. (c) Electrophoresis image of the amplicons. Lane 1 is the first reaction, in which the three fragments were PCR-amplified and subjected to overlap extensions. Lane 2 presents the second reaction, which PCR-amplified the full-length construct from the DNA extracted from the 3-kbp region of the gel.

We used the amplified DNA directly for transfection into amoeba cells, immediately followed by inoculation with APMV. At 1-day post-infection, a part of the cells expressed GFP, indicating that the expression cassette was working upon mimivirus infection (Fig. 2a). From the resultant virus suspension, we first selected neomycin-resistant marker-positive cells, followed by screening GFP-positive wells. We identified five wells exhibiting GFP fluorescence in one 96-well plate. From one of the wells, we successfully isolated the GFP-expressing virus (Fig. 2b). We PCR-amplified the DNA fragments containing marker genes from the recombinant virus (Fig. 2c). These amplified fragments contained the expression cassette and the APMV genomic region that was not included in the donor DNA construct (Fig. 2d), indicating that the expression cassette was integrated into the target locus of the viral genome.

**Figure 2.**
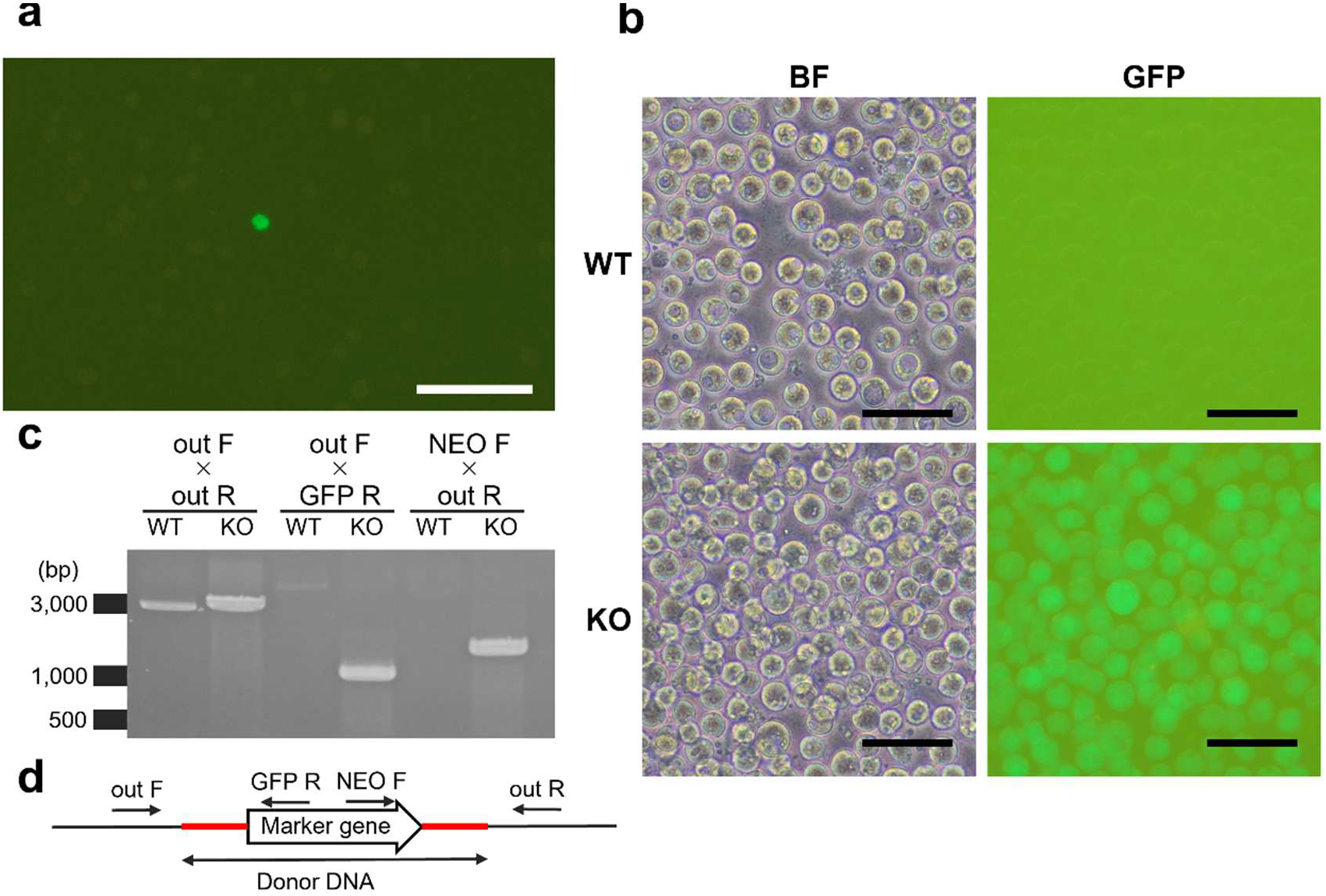
Generation of the recombinant virus. (a) GFP-positive cell found at 1-day post-infection. (b) Amoeba cells infected with the wild-type (WT) and recombinant (KO) APMVs, observed under bright-field (BF) and green fluorescent (GFP). (a, b) Bars, 50 μm. (c) Electrophoresis image for validation of insertion. Each lane represents an amplicon from the WT or KO genomic DNA with the primers designated above. (d) Schematic image of the primer design. The primer sequences are shown in Table 1.

In this study, we developed a method for generating a recombinant mimivirus by directly transfecting a PCR mix into amoeba cells. This method reduces the time and effort required to generate recombinant mimiviruses and could potentially accelerate further reverse-genetics studies focusing on this giant virus. Moreover, our results indicate that PCR-based recombination could be applied to protist viruses. This method retains the potential to be applied to further non-model organisms and viruses.

## Acknowledgements

We thank Prof. Bernard La Scola and Ms. Lina Barrassi for kindly providing APMV. This study was supported by Japan Society for the Promotion of Science KAKENHI grant numbers 21J00174 and 22K15175 to HH and 22H00384 to HO and by Japan Science and Technology Agency ACT-X grant number JPMJAX22BI to HH.

